# Structure insights of the human peroxisomal ABC transporter ALDP

**DOI:** 10.1101/2021.09.24.461756

**Authors:** Yutian Jia, Yanming Zhang, Jianlin Lei, Guanghui Yang

**Affiliations:** State Key Laboratory for Agrobiotechnology, College of Biological Sciences, China Agricultural University, Beijing 100193, China; Technology Center for Protein Sciences, Ministry of Education Key Laboratory of Protein Sciences, School of Life Sciences, Tsinghua University, Beijing 100084, China

## Abstract

Adrenoleukodystrophy protein (ALDP) is responsible for the transport of free very-long-chain fatty acids (VLCFAs) and corresponding CoA-esters across the peroxisomal membrane. ALDP belongs to the ATP-binding cassette sub-family D, which is also named as ABCD1. Dysfunction of ALDP leads to peroxisomal metabolic disorder exemplified by X-linked adrenoleukodystrophy (ALD). Hundreds of ALD-causing mutations are identified on ALDP. However, the pathogenic mechanisms of these mutations are restricted to clinical description due to limited structural information. Furthermore, ALDP plays a role in myelin maintenance, which is tightly associated with axon regeneration. Here we report the cryo-electron microscopy (cryo-EM) structure of human ALDP with nominal resolution of 3.4 Å in nucleotide free state. The structure of ALDP exhibits a typical assembly of ABC transporters. The nucleotide binding domains (NBDs) displays a ligand free state. ALDP exhibits an inward-open conformation to the cytosol. A short helix is located at the peroxisomal side, which is different from other three members of ABCD transporters. The two transmembrane domains (TMDs) of ALDP form a cavity, in which two lipid-like densities can be recognized as the head group of an coenzyme-A ester of a lipid. This structure provides a framework for understanding the working mechanism of ALDP and classification of the disease-causing mutations.

---

Peroxisomes are single-membrane organelles in eukaryotic cells, which play an important role in phospholipid biosynthesis, fatty acid β-oxidation, and hydrogen peroxide degradation (Kao et al., 2018; Wanders and Waterham, 2006). Genetic defects of peroxisomes affect multiple metabolic pathways and lead to severe clinical symptoms, exemplified by X-linked adrenoleukodystrophy (ALD) (Turk et al., 2020). ALD is an inherited disease that mainly affects the nervous system and the adrenal glands in kidney (Lauer et al., 2017; Moser and Hopkins, 1987). X-linked ALD results in aberrant accumulation of VLCFA in the cytosol due to impaired peroxisomal β-oxidation. X-linked ALD leads to progressive demyelination (Berger et al., 2014; van Geel et al., 2001; Rattay et al., 2020; Wiesinger et al., 2013). Loss of myelin reduces the ability of the nerves to transfer information and triggers neuroinflammation. The pathogenesis of ALD is tightly associated by mutations on ALDP (Kemp et al., 1996; Mosser et al., 1993, 1994). Over 900 mutations have been identified on ALDP (https://adrenoleukodystrophy.info/mutations-biochemistry/mutations-biochemistry).

There are other three members of ABCD transporters: ALDP-related protein (ALDRP/ABCD2) (Lombard-Platet et al., 1996), PMP70/ABCD3 (Kamijo et al., 1990), and a cobalamin transporter ABCD4 (Coelho et al., 2012). Among the four members of ABCD transporters, ABCD1-3 are distributed on peroxisomes, while ABCD4 is located on lysosomes. ABCD1-2 are both involved in the transport of VLCFA with different length of acyl tails. ABCD3 is reported to transport long chain fatty acids (LCFA). The orthologues of ABCD transporters in S. cerevisiae contain two half ABC transporters, Pxa1p and Pxa2p, which imports long-chain acyl-CoA esters (Hettema et al., 1996). In contrast to human ABCD1, Pxa1p and Pxa2p function as heterodimers (Hettema et al., 1996). In *Arabidopsis thaliana*, ABCD1 is assembled by a single polypeptide chain but not two subunits (Hurlock et al., 2014; Martinoia et al., 2002; Zolman et al., 2001).

Structural information of ABCD1-3 are eagerly required for the interpretation of lipid transport process in peroxisomes. Investigations on pathological roles of the numerous ALD-causing mutations still await structural elucidation of ALDP. The only structure model of ABCD transporters is ABCD4 bound with ATPs (Xu et al., 2019). However, ABCD4 displays different cellular localization and function from ABCD1-3. ABCD4 specifically transport cobalamin from the lysosome to cytosol (Coelho et al., 2012). In contrast, ABCD1 imports VLCFA from cytosol to peroxisome. Thus, how peroxisomal ABC transporters recognize and import the substrates remains unclear.

In this study, we report the cryo-electron microscopy (cryo-EM) structure of human ALPD/ABCD1 at 3.4 Å resolution in the nucleotides free state. The EM density of ALDP reveals a homo-dimeric assembly. ALDP displays an inward-facing conformation open to the cytosol. We observe a short helix at the peroxisomal side, which is a unique structural element in ABCD transporters. Intriguingly, two lipid-like densities insert into the hydrophobic cavity formed by the two transmembrane domains (TMDs). A head group of a coenzyme-A (Co-A) ester of a lipid fits well in the lipid-like density. Based on the structure, we classified the ALD-mutations into several groups for a better interpretation of their pathogenic roles. The structure of ALDP provides a framework for understanding the mechanism of VLCFA transport from cytosol to peroxisomes.

## Structural elucidation of the human ALDP in nucleotides free state

ABCD1 was expressed in Spodoptera frugiperda cells and purified for cryo-EM analysis (Figure S1). In total 1,922 micrographs were recorded, yielding 1,259,972 particles (Figure S1). The extracted particles were applied to two-dimensional (2D) classifications and three-dimensional (3D) classifications. 289,321 particles were finally used for auto-refinement, yielding a map at an average resolution of 3.4 Å (Figure 1; Figure S1).

**Figure 1.**
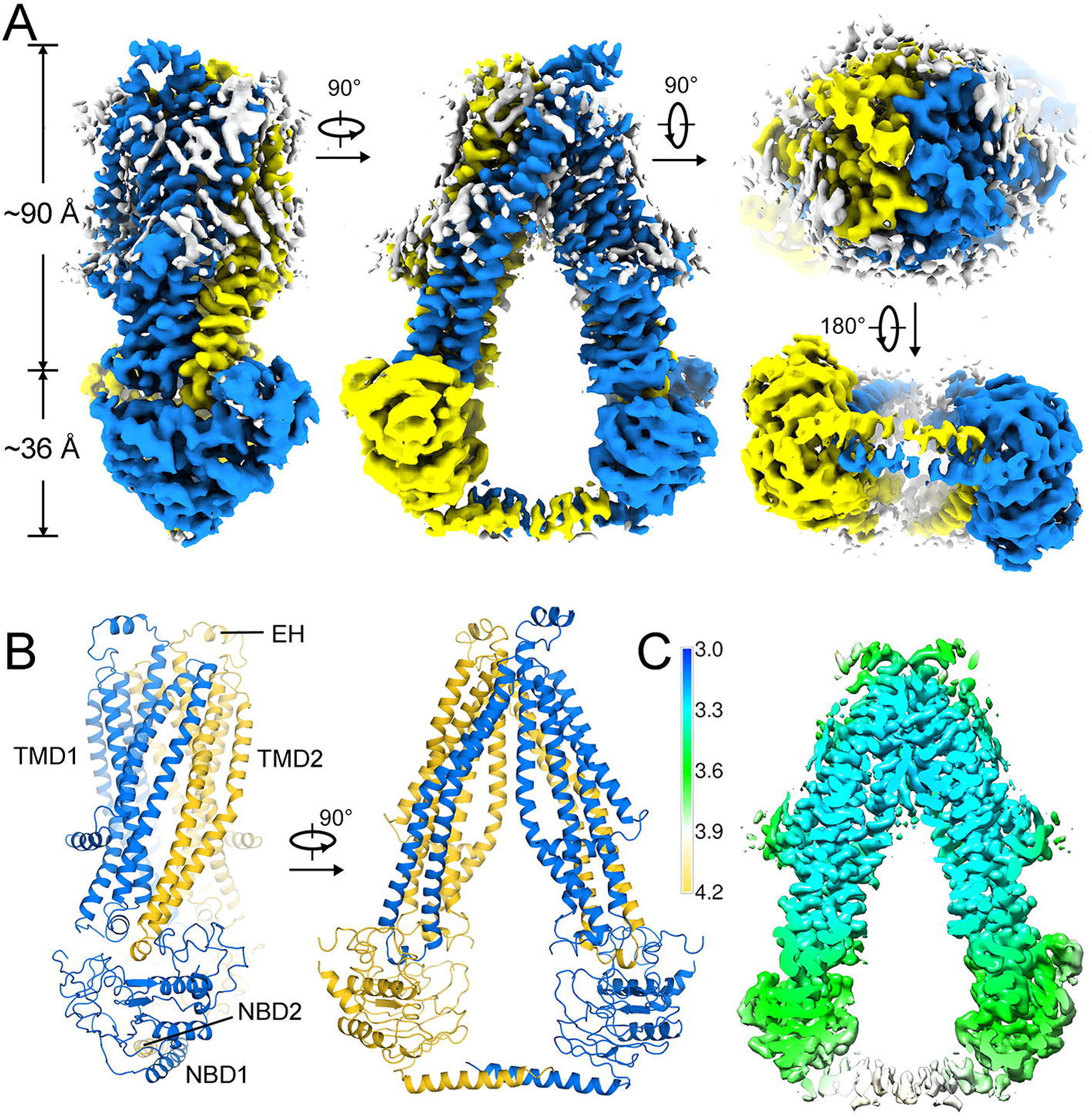
Structure determination of human ALDP. (A) Cryo-EM map of ALDP with each monomer individually colored. The attached lipids around the TMDs are colored in white. (B) The overall structure of ALDP. The TMDs and NBDs are indicated. ALDP contains a short helix at the peroxisomal side. (C) Resolution estimation of the density map.

The model of ABCD1 were *de novo* built for each subunit (Figure 1, Figure S2). The TMD of each subunit contains six transmembrane helices, which exhibit a domain-swapped arrangement. The TM4-5 and their intervening helix of one subunit are coordinated by TM2-3 and NBD of the other subunit (Figure 1 and 2). In the absence of nucleotides, the two NBDs at the cytosolic side separate with each other about 40 Å. A short helix located at the peroxisomal side can be unambiguous identified, which is a featured structural element of ALDP. We observed two extended helical-shaped densities at the carboxyl-terminus of NBDs. However, the local resolution of these densities are relatively poor at about 4.0 Å, which is not sufficient to build an accurate model. We analyzed the predicted model by Alphafold (Jumper et al., 2021), which has a C-terminal helix predicted in high confidence (Figure S3). Thus, we putatively docked two short helices in the C-terminal densities of ABCD1.

**Figure 2.**
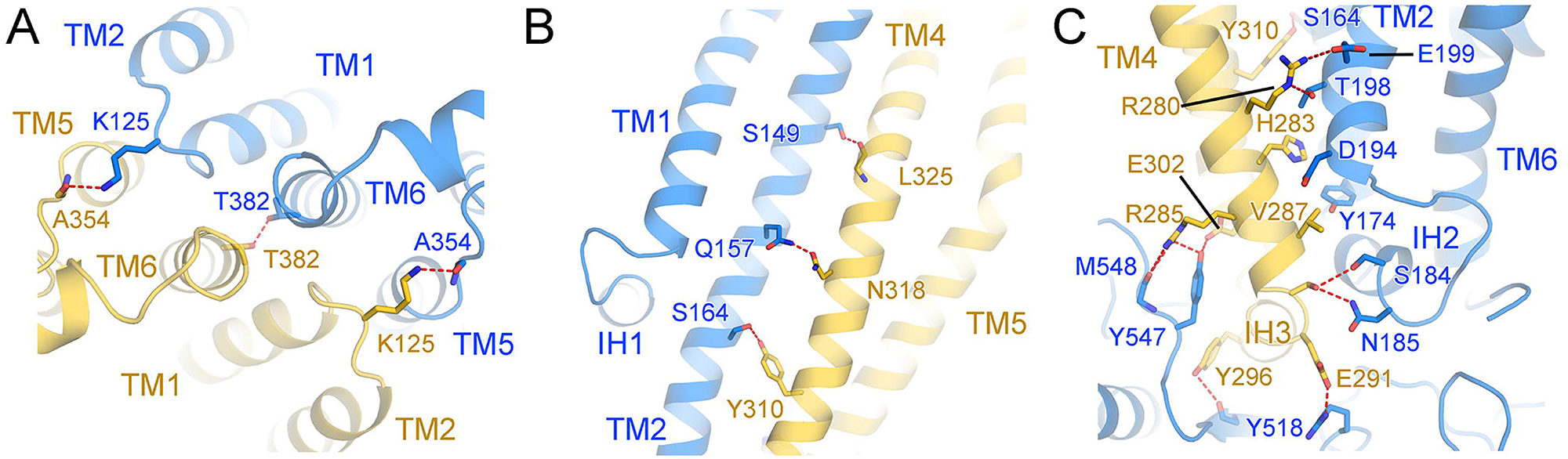
Dimeric assembly of ALDP. (A-B) Two subunits of ALDP stack against with each other to form a homodimer. The dimeric assembly is stabilized by hydrogen bonds between the TMs. The interactions between the TM4-5 and the opposing NBD shown in (C).

Two lipid-like densities can be found to insert the cavity formed by the two TMDs. These two densities can be clearly discriminated from the other lipid densities surrounding the TMDs. The head group of a Co-A ester can fit well into the two densities (Figure 3). These two lipids seem to be endogenous lipids that co-purified with ALDP because no exogenous lipids were added during purification. Nonetheless, the Co-A ester form of VLCFA is the substrate of ALDP. We putatively built two Co-A esters into these densities (Figure 3). The residues surrounding the putative Co-A esters are all found to be the sites for ALD-mutations (Figure 3C). Such observations confirms the substrate binding site of ALDP.

**Figure 3.**
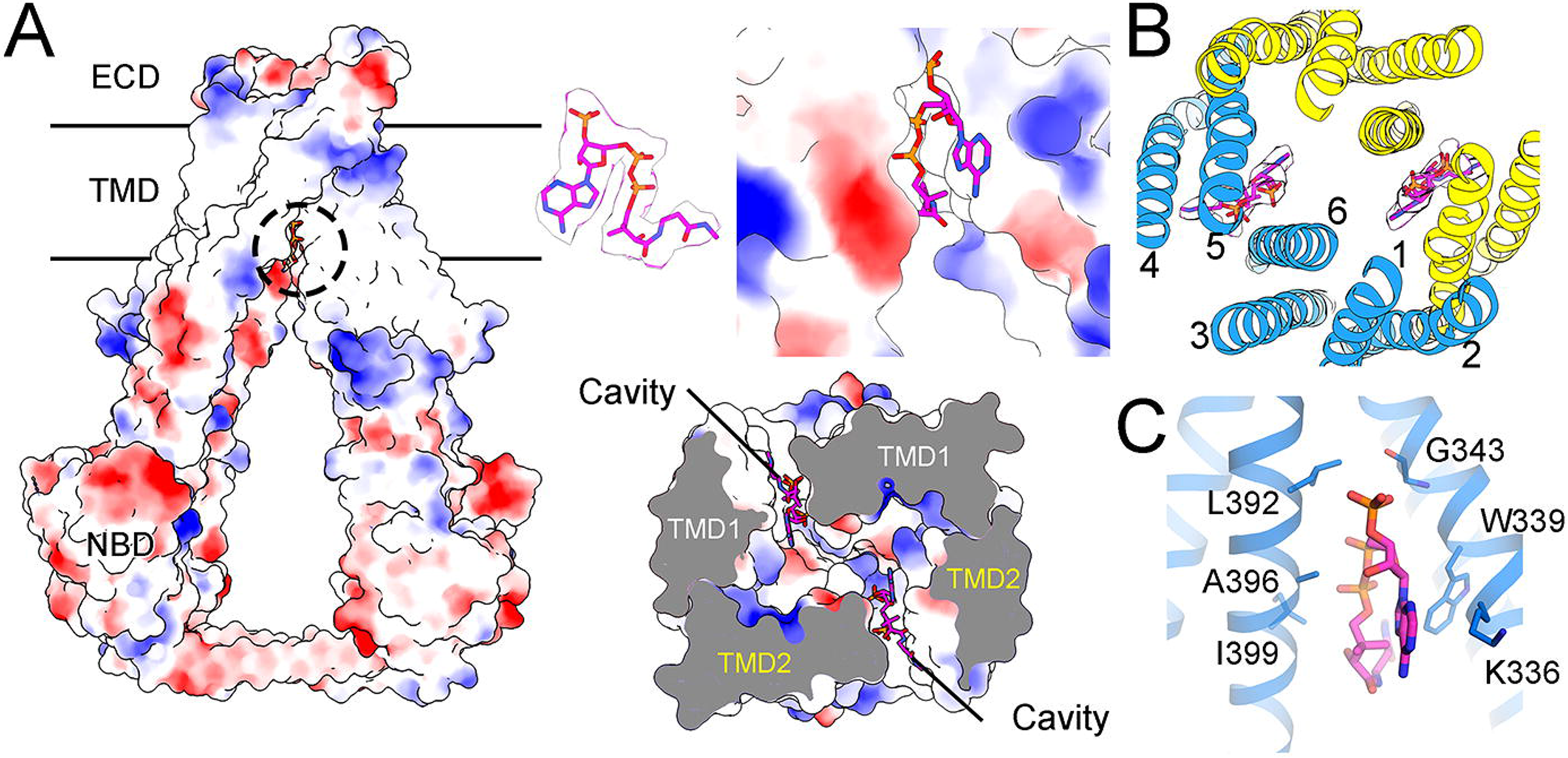
Coordination of two putative Co-A esters by ALDP. (A) The surface electrostatic potential of ALDP (left). The putative Co-A esters insert into the cavity formed by TMDs (right). (B) Coordination of Co-A esters by each protomer of ALDP. The residues surrounding the Co-A ester are shown in (C). All these residues are reported to be mutation sites in ALD..

**Figure 4.**
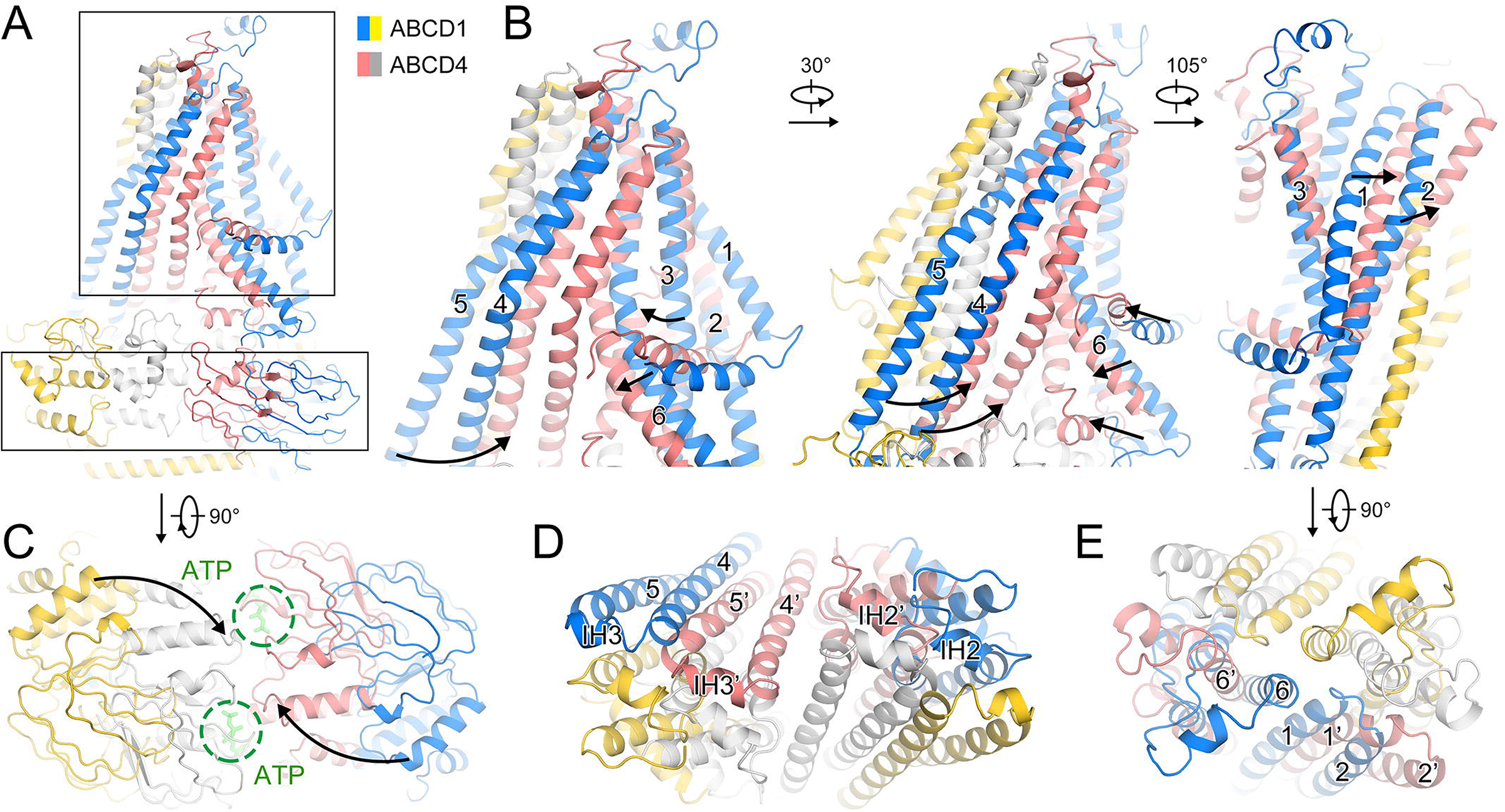
Distinct conformations between ALDP and ATP-loaded ABCD4. (A) Structural comparison of ALDP and ABCD4. ALDP displays a nucleotide free conformation. The Glu549 of ABCD4 is substituted by Gln to avoid ATP hydrolysis. (B) Superimposition of the TMDs of ATP-free ALDP and ATP-bound ABCD4. The TM4-5 and TM6 of ABCD4 undergo large conformational changes towards the cavity, resulting a narrow cavity than that of ALDP. (C) The bottom view of the NBDs of ATP-free ALDP and ATP-bound ABCD4. (D-E) The cut view of the TMDs from the cytosolic and the peroxisomal side.

**Figure 5.**
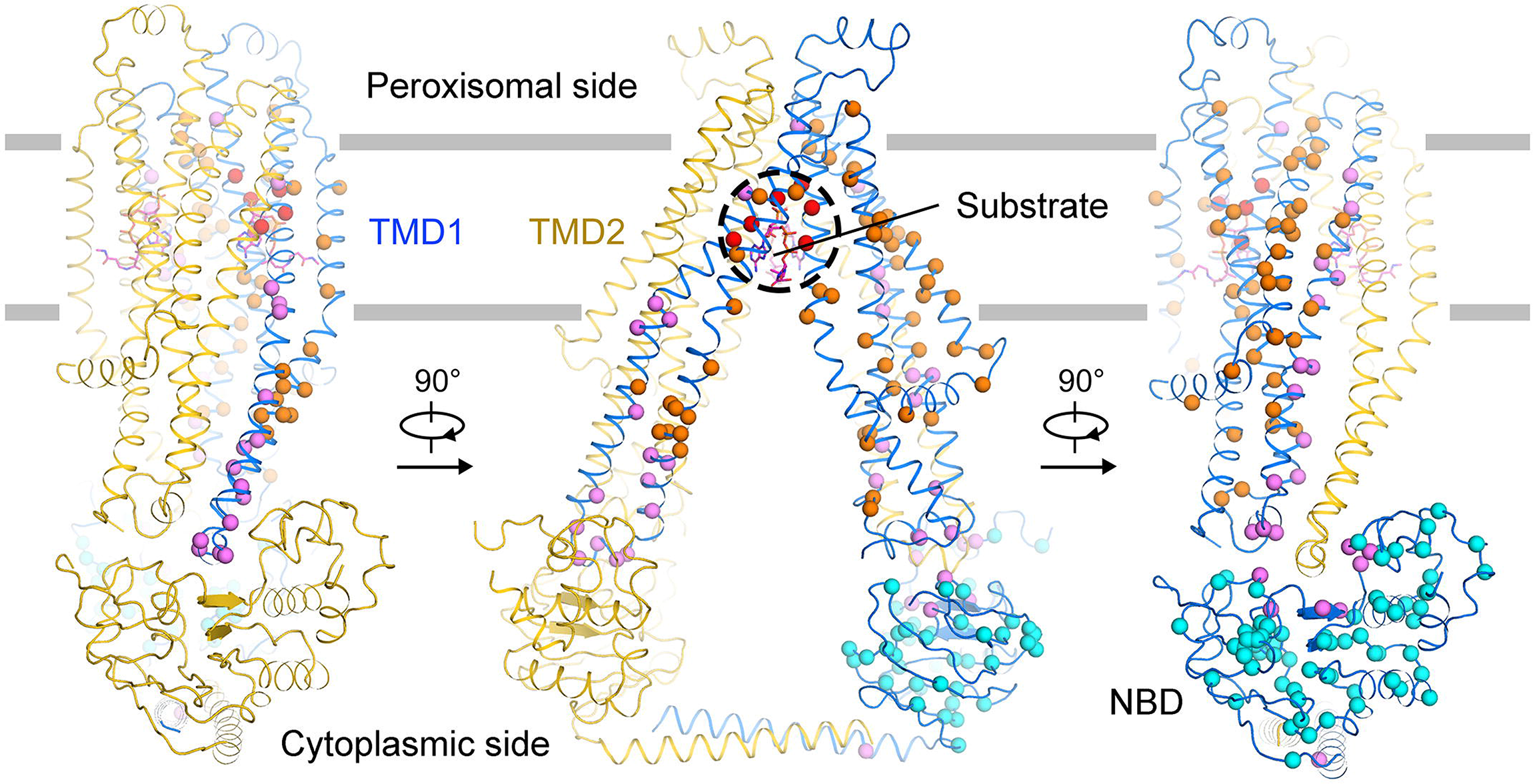
Structural based classification of the ALD-derived mutations. Structural mapping of clinical-derived pathogenic mutations of ALDP. The Cα atoms of the indicated residues are shown as spheres. The residues surrounding the putative CoA-ester group at the lipid cavity shown as red spheres. The residues which mediate the dimeric assembly of ALDP are colored in violet. Other residues on TMD and NBD are presented in orange and cyan, respectively.

## Distinct conformations between ALDP without nucleotide and ABCD4 bound with ATP

Although ALDP and ABCD4 shares 22% sequence identity, the overall structural organization of the TMD and NBD are similar except for the peroxisomal helix and the putative C-terminal helix. In the absence of nucleotides, ALDP displays an inward open conformation to the cytosol. ABCD4 exhibits a lysosome-open conformation bound with ATP (Xu et al., 2019). We aligned the structures of ALDP and ABCD4. In the presence of ATP, the TM4-5 and TM6 of ABCD4 move forward to each other, resulting in a narrow cavity than that of ALDP. Such conformational changes are triggered by binding of nucleotides. Upon binding of ATP, the NBDs of ALDP may undergo large conformational changes, accompanied with the movement of TM4-5 of the neighboring subunit. Conformational changes of the TM4-5 may lead to the movement of TM2 through reorganizing the hydrogen bonds between TM4 and TM2 (Figure 2). The TM2 serves as a mechanical lever: the cytosolic side moves towards the central cavity, while the peroxisomal side may move backwards the symmetry axis of the dimer. Such concerted conformational changes allow the delivery of substrates across the organelle membranes.

To elucidate the transport mechanism of the ABCD transporters, we compared the structures of all four members (Figure S4). Structures of ALDP and ABCD4 are determined by cryo-EM, while the structures of ABCD2 and ABCD3 are predicted by Alphafold (Jumper et al., 2021). The TMDs and NBDs of all four members exhibit high similarity. These four structures vary mostly at the C-terminus. Based on the sequence alignment, ABCD1-2 harbor more residues at the C-terminus that form a helix. Although the function of ABCD3 in transport LFCA is relevant to ABCD1-2, it shares higher sequence and structural similarity with ABCD4. The structural arrangement of ALDP is reminiscent of several other ABC transporters as well (Figure S5).

## Structural based classification of pathogenic mutations on ALDP

Previous biochemical studies have been carried out to reveal the pathogenic mechanisms of these ALD-associated mutations (Braun et al., 1995; Fanen et al., 1997; Fuchs et al., 1994; Gartner et al., 1997; Imamura et al., 1997; Ligtenberg et al., 1995). Whether the plenty of mutations affect the transporter in similar manner remains to be investigated. However, individually examination of these mutations requires enormous and redundant efforts. For a better interpretation of the mutations, a brief classification of these mutations into several groups may facilitate the functional studies. Our structure provides such a framework to have these mutations classified into several groups. The first group is the residues that located at the substrate binding cavity. We have identified the residues that may be directly involved in substrate binding (Figure 2). The second group of residues are mainly distributed at the dimer interfaces of the two subunits. Pathogenic mutations on these residues may disturb the stability of ALDP. The third group are located on the NBDs, which may hinder the hydrolysis of nucleotides. The last group are mainly mapped to the TMDs, which may affect the conformational changes during substrate transport. It should be noted that there are several residues belong to one more groups. For example, the residues on the NBD that coordinate the TM4-5 can be classified into group II and III.

## Discussion

The structure of ALDP provides an important framework to illustrate the molecular basis of VLCFA transport to peroxisomes and the pathogenic roles of ALD-derived mutations. At the current stage, we observe the nucleotide-free conformation of ALDP, which opens to the cytosol and close at the peroxisomal side. Two endogenous lipid can be clearly identified at the 3.4 Å map, which insert into the substrate binding cavity. The head group can be recognized as a Co-A ester, while the long hydrophobic tail of this lipid is absent due to high flexibility. Structural comparison of ALDP to other ABC transporters reveals the possible conformational changes during a whole cycle of substrate transport. Further studies on ALDP may focus on the peroxisomal open state in the presence of nucleotides and biochemical characterizations of the ALD-mutants based on the structures.

## ACKNOWLEDGEMENTS

We thank the Tsinghua University Branch of China National Center for Protein Sciences (Beijing) for the cryo-EM facility and the computational facility support, Dr. Xiaomin Li, Dr. Fan Yang and Tao Liu for technical support in EM data acquisition. This work was supported by National Natural Science Foundation of China (3217110084); Chinese Universities Scientific Fund funds (15050004, 15050017, 15051002) and Young Elite Scientists Sponsorship Program by China Association for Science and Technology.

## DATA AND SOFTWARE AVAILABILITY

The cryo-EM map of ALDP has been deposited in the Electron Microscopy Data Bank (EMDB) with the accession code EMD-XXXXX. The atomic coordinates for the corresponding model have been deposited in the Protein Data Bank (PDB) under the accession code XXXX.

