## Supplemental Figures for "Structure insights of the human peroxisomal ABC transporter ALDP"

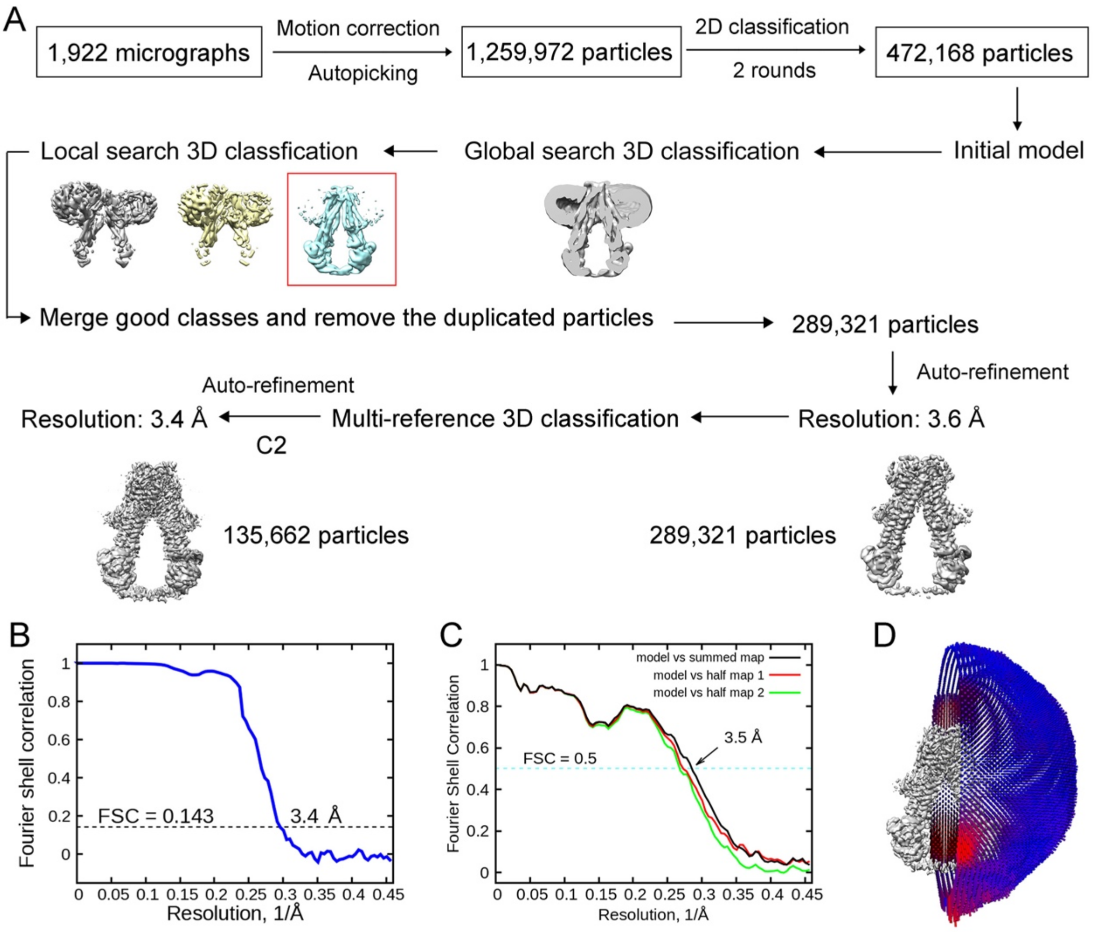


**Figure S1. Cryo-EM analysis of ALDP.**

(A) A flowchart description of EM data processing. (B) The FSC curve for the cryo-EM map. The average resolution is estimated to be 3.4 Å on the basis of the FSC value of 0.143. (C) FSC curves of the refined model versus the overall 3.4-Å map that it was refined against (black); of the model refined in the first of the two independent maps used for the FSC calculation versus that same map (red); and of the model refined in the first of the two independent maps versus the second independent map (green). The small difference between the red and green curves indicates that the refinement of the atomic coordinates did not suffer from overfitting. (D) Angular distribution of the particles used for reconstruction of the ALDP. Each cylinder represents one view and the height of the cylinder is proportional to the number of particles for that view.

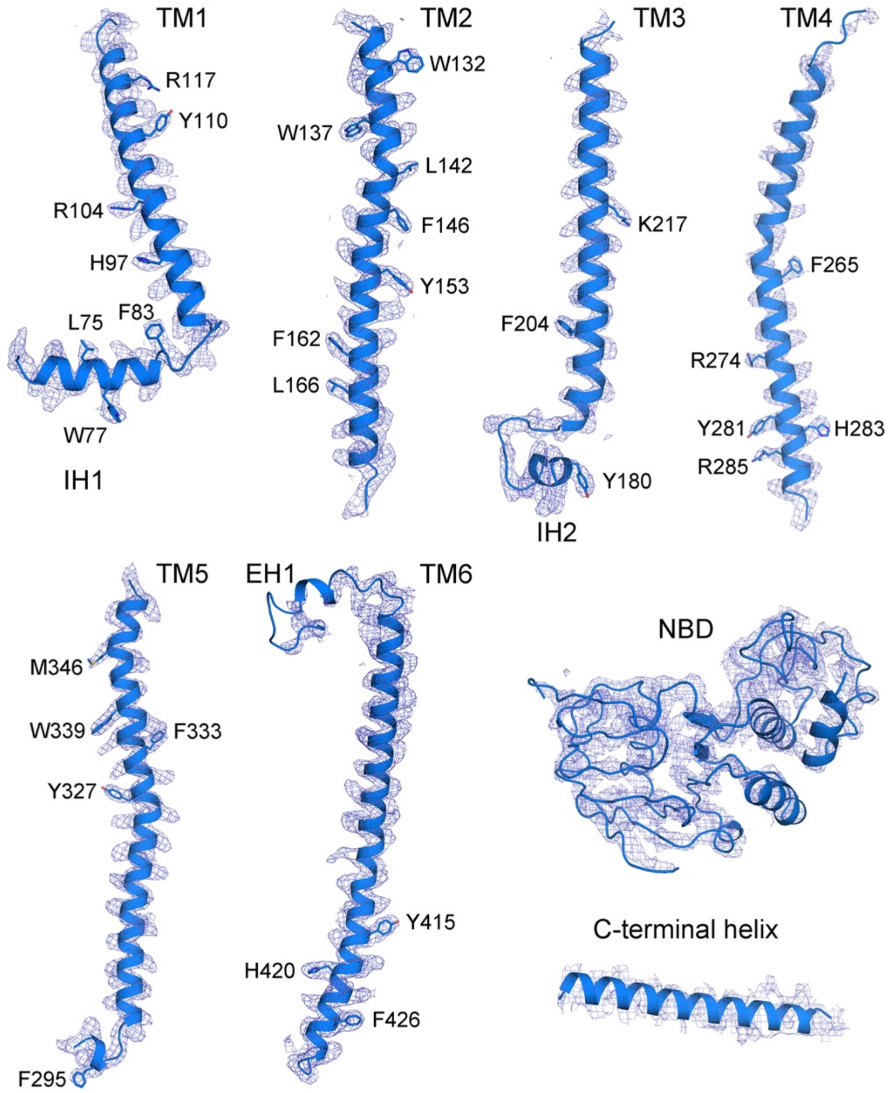


**Figure S2. Representative cryo-EM densities of structural elements of ALDP.**

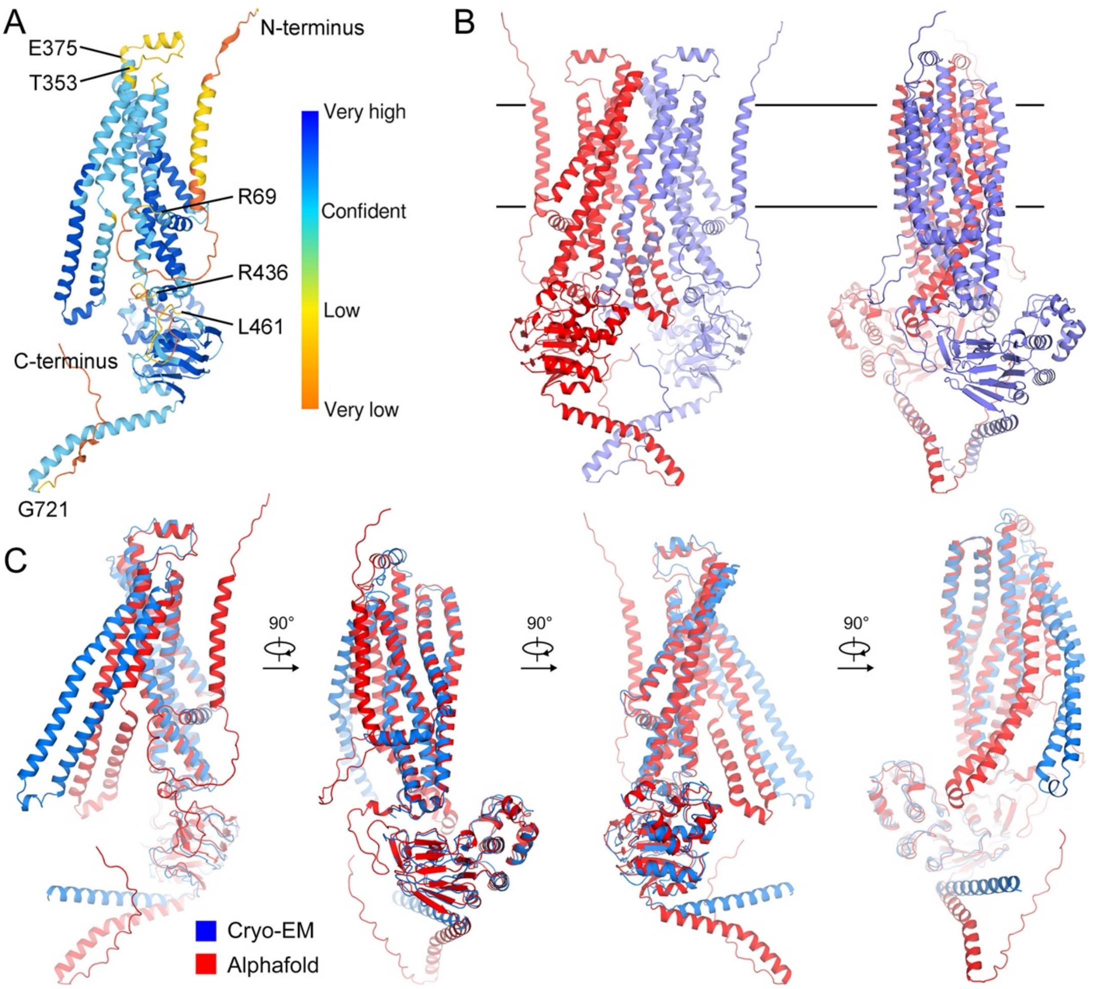


**Figure S3. Structural comparison between cryo-EM structure and the predicted one by Alphafold.**

(A) A predicted model of ALDP by AlphaFold (AF-ALDP). (B) The dimeric structural model of AF-ALDP generated through structural superimposition to the cryo-EM structure. (C) Structural comparison between the predicted model and the cryo-EM model.

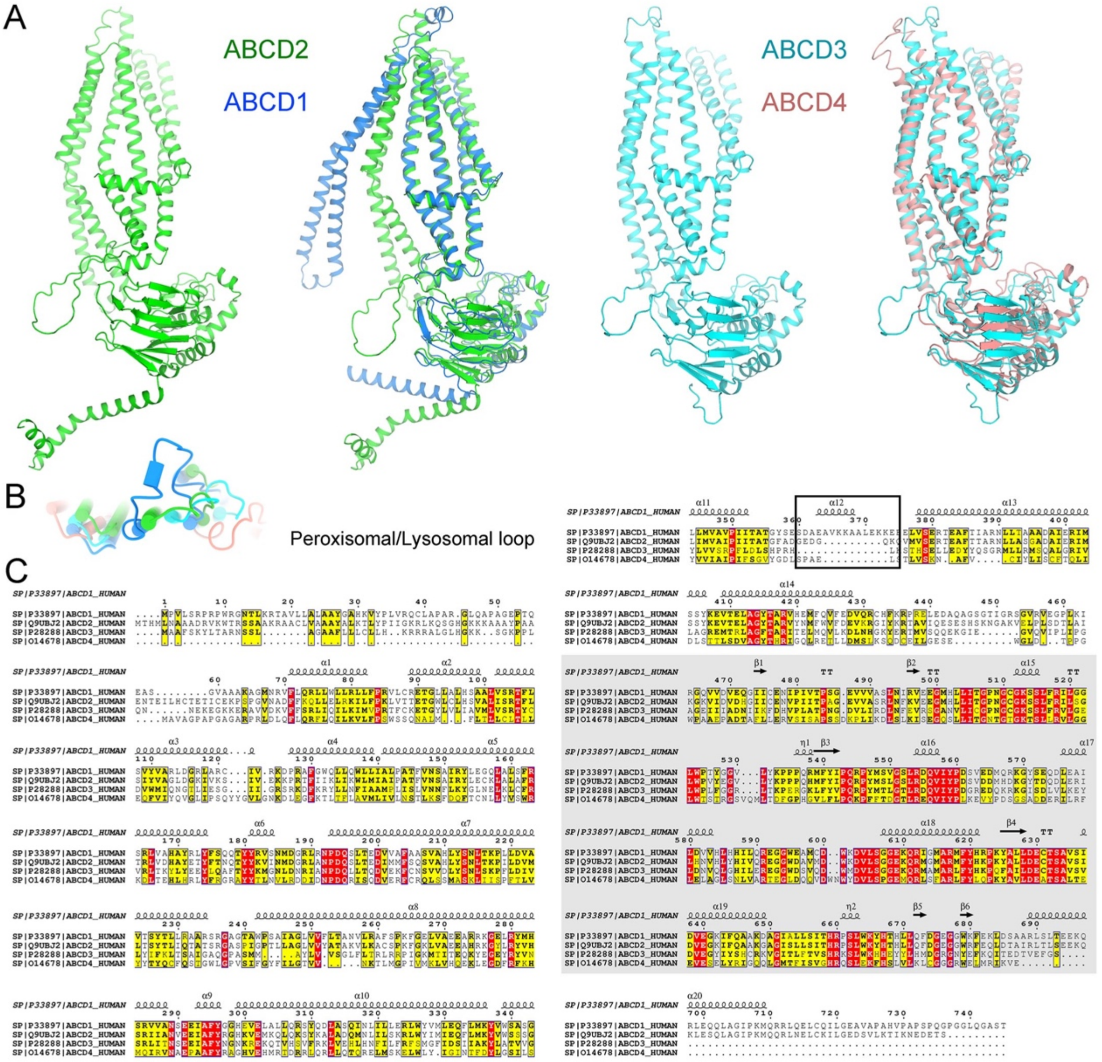


**Figure S4. Structural comparison of all members of ABCD transporters in human.** (A) Structure similarities of the ABCD members. High structural similarities are found between ALDP and ABCD2 or ABCD3 and ABCD4. The model of ABCD2 and ABCD3 are predicted by Alphafold. (B) Distinctive extracellular helix of ALDP at the peroxisomal side. (C) Sequence alignment of ABCD members. The sequences corresponding to the short peroxisomal helix are highlighted.

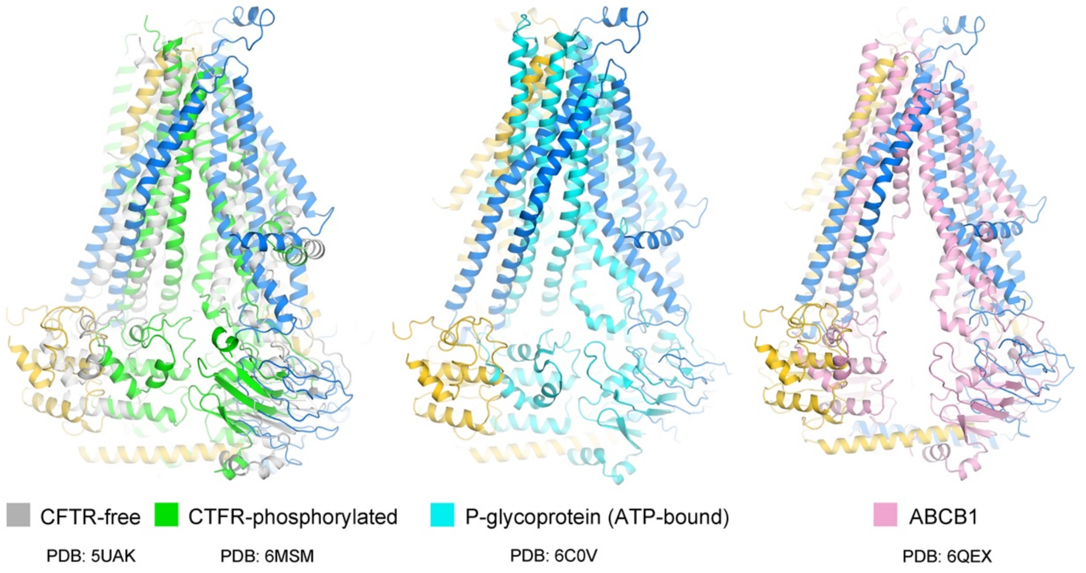


**Figure S5. Structural alignment of ALDP to other representative ABC transporters.**
